# High Angular Resolution Susceptibility Imaging and Estimation of Fiber Orientation Distribution Functions in Primate Brain

**DOI:** 10.1101/2022.10.23.513390

**Authors:** Dimitrios G. Gkotsoulias, Roland Müller, Carsten Jäger, Torsten Schlumm, Toralf Mildner, Cornelius Eichner, André Pampel, Jennifer Jaffe, Tobias Gräßle, Niklas Alsleben, Jingjia Chen, Catherine Crockford, Roman Wittig, Chunlei Liu, Harald E. Möller

**Affiliations:** Nuclear Magnetic Resonance Methods & Development Group, Max Planck Institute for Human Cognitive and Brain Sciences, Leipzig, Germany; Department of Neurophysics, Max Planck Institute for Human Cognitive and Brain Sciences, Leipzig, Germany; Department of Neuropsychology, Max Planck Institute for Human Cognitive and Brain Sciences, Leipzig, Germany; Max Planck Institute for Evolutionary Anthropology, Leipzig, Germany; Taï Chimpanzee Project, Centre Suisse de Recherches Scientifiques en Côte d’Ivoire, Côte d’Ivoire; Helmholtz Institute for One Health, Greifswald, Germany; Robert Koch Institute, Epidemiology of Highly Pathogenic Microorganisms, Berlin, Germany; Electrical Engineering and Computer Sciences, University of California, Berkeley, CA, USA; Institute of Cognitive Sciences, CNRS UMR5229 University of Lyon, Bron, France; Helen Wills Neuroscience Institute, University of California, Berkeley, CA, USA

**Keywords:** Anisotropic magnetic susceptibility, diffusion-weighted imaging, gradient-recalled echo, high angular resolution, orientation distribution function, quantitative susceptibility mapping

## Abstract

Uncovering brain-tissue microstructure including axonal characteristics is a major neuroimaging research focus. Within this scope, anisotropic properties of magnetic susceptibility in white matter have been successfully employed to estimate primary axonal trajectories using mono-tensorial models. However, anisotropic susceptibility has not yet been considered for modeling more complex fiber structures within a voxel, such as intersecting bundles, or an estimation of orientation distribution functions (ODFs). This information is routinely obtained by high angular resolution *diffusion* imaging (HARDI) techniques. In applications to fixed tissue, however, diffusion-weighted imaging suffers from an inherently low signal-to-noise ratio and limited spatial resolution, leading to high demands on the performance of the gradient system in order to mitigate these limitations. In the current work, high angular resolution *susceptibility* imaging (HARSI) is proposed as a novel, phase-based methodology to estimate ODFs. A multiple gradient-echo dataset was acquired in an entire fixed chimpanzee brain at 61 orientations by reorienting the specimen in the magnetic field. The constant solid angle method was adapted for estimating phase-based ODFs. HARDI data were also acquired for comparison. HARSI yielded information on whole-brain fiber architecture, including identification of peaks of multiple bundles that resembled features of the HARDI results. Distinct differences between both methods suggest that susceptibility properties may offer complementary microstructural information. These proof-of-concept results indicate a potential to study the axonal organization in *post-mortem* primate and human brain at high resolution.

**Highlights:**

- Introduction of High Angular Resolution Susceptibility Imaging (HARSI) for advancing Quantitative Susceptibility Mapping (QSM).
- HARSI-derived fiber orientation distributions in fixed chimpanzee brain.
- HARSI-based visualization of complex fiber configurations.
- Comparisons between HARSI and High Angular Resolution Diffusion Imaging.
- Potential for high-resolution *post-mortem* imaging of fiber architecture.

## 1. Introduction

Volume magnetic susceptibility (χ) is a dimensionless quantity (in SI units), which describes the degree of magnetization of a material placed inside an external magnetic field. In biological tissues, local structural characteristics lead to spatial variations of susceptibility and, thus, to local field disturbances and measurable differences in the resonance frequency in magnetic resonance imaging (MRI). Based on the nature of these disturbances, susceptibility may be paramagnetic (χ > 0) or diamagnetic (χ < 0). In reverse, quantifying the susceptibility properties of tissue can lead to a non-invasive elucidation of microstructural properties and details at the whole-brain scale, without the need for sophisticated microscopy techniques (Möller et al., 2019).

Quantitative susceptibility mapping (QSM) methods provide voxel-wise susceptibility estimations based on the signal phase in gradient-recalled echo (GRE) acquisitions and a series of post-processing steps (Deistung et al., 2017; de Rochefort et al., 2010; Liu et al., 2015; Shmueli et al., 2009). Although various approaches have been developed for this purpose, robust mapping of χ from MRI phase data remains challenging because the field perturbation (and, hence, the signal phase) associated with a susceptibility distribution is inherently non-local and depends on the distribution’s orientation in the magnet (Schäfer et al., 2009). Briefly, following appropriate combination of the complex multi-channel data from a phased-array coil and removal of phase wraps (Robinson et al., 2017) and background-field contributions (Schweser et al., 2017), field-to-source inversion methods are used to solve (in the Fourier domain) the ill-posed problem of going from the signal phase (as a measure of the local magnetic flux density offset, *δB*_0_) to the local variations in χ. Notably, this is typically performed under the assumption that χ representing an imaging voxel is a scalar, isotropic quantity (Deistung et al., 2017). While this assumption may be justified in regions of cerebral gray matter (GM), multiple studies have shown that the bulk susceptibility in cerebral white matter (WM) exhibits a highly anisotropic character—primarily, as a result of the specific arrangement of the myelin sheaths enveloping axons at a molecular level, leading to an accumulated macroscopic effect on the voxel level (Li et al., 2017; Liu, 2010; Wharton & Bowtell, 2012).

Consequently, brain tissue susceptibility cannot be fully understood and exploited if its anisotropic characteristics are not taken into consideration—similarly to diffusion tensor imaging (DTI) that exploits the anisotropic characteristics of diffusion as an expansion of simple measurements of the mean diffusivity (MD). Methods proposed for addressing this issue, such as susceptibility tensor imaging (STI), consider multiple GRE acquisitions by reorienting the object (e.g., the human head) inside the magnet—as current MRI technology does not allow to reorient the magnetic field around the imaged object (Liu, 2010). In the basic STI approach, susceptibility is depicted as a second-rank symmetric tensor with six unique parameters, in a similar way as orientation-dependent diffusivity is modeled in DTI (Basser et al., 1994a; 1994b). Obtaining GRE phase measurements along six or more unique orientations allows for the reconstruction of a second-rank tensor χ, based on a linear system of equations constructed by the relationship (Li et al., 2017; Liu, 2010):

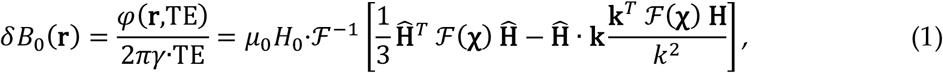

In Eq. 1, *φ*(**r**,TE) is the signal phase in image space at position **r** and echo time TE, *γ* is the gyromagnetic ratio, µ_0_ is the vacuum permeability, *H*_0_ and 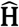 are, respectively, the magnitude of the applied magnetic field and its unit vector (defining the laboratory frame’s *z*-direction), and **k** is the spatial frequency vector. ℱ and ℱ^−1^ denote the Fourier transform and its inverse, respectively, and the subscript ^*T*^ the transpose. Note that the χ_33_ tensor component has also been suggested as an STI-based estimation of a scalar susceptibility (Langkammer et al., 2018).

Despite being a useful approximation of the diffusion signal for a certain range of *b*-values (Novikov et al., 2018), DTI is mathematically incapable of resolving multiple fiber orientations within a voxel, thus providing limited information for the vast majority of WM regions (Jones et al., 2013). Important improvements are achieved with High Angular Resolution Diffusion Imaging (HARDI) techniques, sampling 60 or even more diffusion directions (Frank et al., 2001; Tournier et al., 2004; Tuch et al., 2002; 1999). This includes high-quality estimations of fiber orientation distribution functions (ODFs) and resolving intersecting fiber bundles. Compared to these developments in diffusion-weighted imaging (DWI), the need for physical rotation of the object and the complex processing pipeline of multi-orientational susceptibility imaging has led to datasets of limited angular resolution so far. For example, STI acquisitions in formalin-fixed mouse brain specimens were obtained with 19 orientations, which were evenly distributed on a spherical surface if a sufficiently large radiofrequency (RF) coil could be used (Liu et al., 2012) but restricted to rotations about the sample’s long axis with a tightly fitting solenoid (Li et al., 2012). *In-vivo* acquisitions in humans achieved 16 rotation angles (about the anterior-posterior and the left-right direction) up to ±50° with a relatively open quadrature coil (Li et al., 2012) but only 12 orientations with angles up to ±25° with a spatially more constrained head array (Bilgic et al., 2016a). Such constraints impose limitations on the quality of tensor-based analyses and, more importantly, prohibit to go beyond STI for microstructural information, such as susceptibility-based ODFs (Liu et al., 2013).

In the current work, we introduce High Angular Resolution Susceptibility Imaging (HARSI) as an advanced QSM approach for *post-mortem* acquisitions—similar to HARDI techniques developed in the context of DWI. The goal is to investigate the orientation-dependent susceptibility at high angular resolution to achieve WM characterization beyond state-of-the-art tensor-based methods. A multi-echo (ME) GRE dataset comprising 61 unique directions is presented, and HARSI-based STI estimates are compared to single-orientation QSM results. Additionally, ODFs are obtained by applying the generalized constant solid angle (CSA) method (Kamath et al., 2012). The results indicate comparable potential in resolving intersecting fiber orientations as HARDI-based ODFs and suggest strong prospects for obtaining complementary information on WM microstructure.

## 2. Methods

### 2.1. Brain specimen

The *post-mortem* specimen used for the acquisitions was a whole brain obtained from an adult wild alpha-male chimpanzee (*Pan troglodytes verus*; male, 45 years). The animal had died from natural causes in Taï National Park (Parc National de Taï), Côte d’Ivoire, without human interference. Approximately 18 h after death, a specifically trained veterinarian performed the brain extraction wearing full personal protective equipment and adhering to the necroscopy protocols at the field site. All procedures followed the ethical guidelines of primatological research at the Max Planck Institute for Evolutionary Anthropology, Leipzig, which were approved by the Ethics Committee of the Max Planck Society. Immediately after extraction, the brain was preserved by immersion in 4% paraformaldehyde (PFA). The specimen was transferred to Germany under strict observation of CITES (Convention on International Trade in Endangered Species of Wild Fauna and Flora) protocol regulations. After fixation for 6 months, superficial blood vessels were removed, and the PFA was washed out in phosphate-buffered saline (PBS) at pH 7.4 for 24 days.

### 2.2. Brain container for reorientation imaging

For preparatory experiments, the specimen was centered in an oval-shaped acrylic container of suitable size (15cm long-axis and 10cm short-axis diameter) and stabilized with sponges. The container was filled with liquid perfluoropolyether (Fomblin^®^; Solvay Solexis, Bollate, Italy) to protect the tissue from dehydration and to achieve approximate matching of the susceptibility at the brain surface (Benveniste et al., 1999). With this setup, a three-dimensional (3D) high-resolution *T*_1_-weighted MP2RAGE (Magnetization-Prepared 2 RApid Gradient Echoes) dataset was acquired.

Phase-sensitive acquisitions at multiple orientations with respect to the main magnetic field require physical rotations of the object. During these measurements, the sample may be subjected to gravity-induced non-linear deformations, which are inconsistent between scans unless measures are taken to preserve the shape. This leads to inaccuracies and may introduce artifacts during post-processing, which requires excellent registration of the acquired volumes. In previous *post-mortem* experiments in mice (Li et al., 2012; Liu et al., 2012) and also in humans (Alkemade et al., 2020; 2022), the problem of inconsistent deformations was avoided by keeping the fixed brain within the skull after surgical separation of the head. In the current work, a specific brain mask was derived from the 3D *T*_1_-weighted dataset and a custom-made container was designed consisting of an inner and an outer part (Figure 1B–E). Based on the mask, the surface of the individual anatomy was reconstructed and split into a top and a bottom mesh using Python (Figure 1B). The algorithm further allowed for a parameterization of the design, for example, to consider an additional distance from the tissue or modify the wall thickness for sufficient stability. Stereolithography (STL) files were then produced using CAD (computer-aided design) software (Fusion 360^®^; Autodesk, San Rafael, CA, USA). The outer container was a spherical structure of sufficient size (16cm diameter) to take up the anatomically shaped inner container, which was rigidly connected via eight adjustable screws (Figure 1E). The spherical outer shell also consisted of two parts (top and bottom) with further indications of 60 unique orientations on its surface. The orientations were calculated employing an electrostatic repulsion optimization model using *MRtrix3* (Jones et al., 1999; Tournier et al., 2019). The outer container was then positioned on a custom-made holder that included an additional position indicator (Figure 1E). The combined setup ensured robust positioning of the specimen in the RF coil with an orientation error ≤3° for all axes. Note that a refined orientation information was obtained during post-processing from image registration (see below) and used in all subsequent analyses. All container and holder parts were 3D-printed on an Objet Eden260VS (Stratasys, Eden Prairie, MN, USA) using Objet MED610 Biocompatible Clear material (Stratasys).

**Figure 1.**
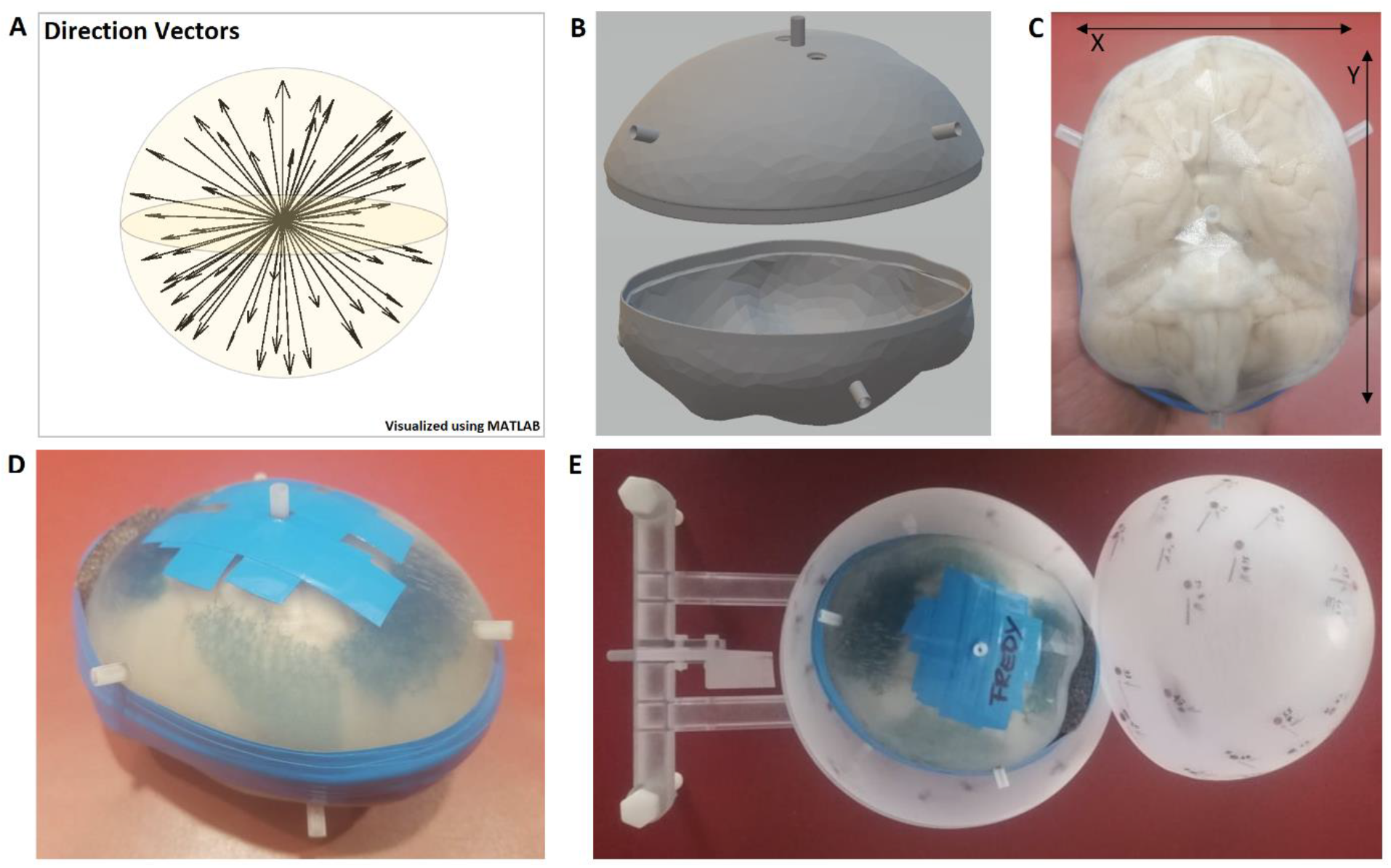
**(A)** Direction unit vectors of 61 evenly distributed orientations on the surface of a sphere as calculated by *MRtrix3*. **(B)** Design of the inner container based on the surface rendering of a 3D *T*_1_-weighted dataset. **(C, D)** 3D-printed container, adapted to the brain’s individual anatomy with indications of the approximate dimensions. The specimen is immersed in Fomblin, and the container is carefully sealed using tape (*x*=105mm, *y*=125mm). **(E)** View of the specimen “Fredy” inside the anatomical container, positioned inside an outer, spherical container with indications of the calculated directions for reorientations. The outer container is positioned on a custom-made holder with an additional angle indicator that supports robust positioning in the RF coil and accurate reorientation.

### 2.3. Image acquisition

All MRI experiments were performed at 3 T on a MAGNETOM Skyra Connectom A (Siemens Healthineers, Erlangen, Germany) that achieves a maximum gradient strength of 300 mT/m (Fan et al., 2022). In order to avoid damage to the gradient coil during long scanning sessions, a first-order approximation of the acoustic response was computed in Matlab from the simulated gradient time course (IDEA DSV file) of each individual imaging protocol with its specific acquisition parameters (Labadie et al., 2013) and carefully inspected for potentially harmful vibrations. The brain container and holder were then positioned inside a 32-channel receive array coil (Siemens Healthineers), and a complex-valued 3D Fast Low-Angle SHot (FLASH) reference dataset (Frahm et al., 1986) was acquired after shimming. Subsequently, 60 additional consecutive acquisitions were performed with varying sample orientations and the same parameters as in the reference scan (1mm isotropic nominal resolution; field of view, FOV = 160×160×160 mm; RF pulse flip angle, *α* = 30°; repetition time, TR = 50 ms; 12 echoes with TE = 3.54 ms and 6.98 ms for the first two echoes and an echo spacing, ΔTE = 3.75 ms, for the remaining 10 echoes; bandwidth, BW = 600 Hz/pixel). GeneRalized Autocalibrating Partially Parallel Acquisitions (GRAPPA) with acceleration factor *R* = 2 (Griswold et al., 2002) and a partial-Fourier scheme (Feinberg et al., 1986) with partial-Fourier factor, *f*_*p*_ = 7/8 were employed in phase-encoding (PE) direction (always along the physical *z*-axis), to accelerate the measurements (acquisition time, TA ≈ 12 min per orientation; total TA ≈ 12:12 h). The slab orientation was always along the *y*-axis, and the vendor’s 3D distortion correction was applied to all acquisitions.

Additional DWI data were acquired at 1mm isotropic nominal resolution employing a previously described 3D segmented ME echo planar imaging (EPI) technique optimized for PFA-fixed tissue (Eichner et al., 2020). Briefly, the sequence consists of a Stejskal-Tanner sequence with multiple gradient-echo refocusing of the first (spin) echo. Combination of the GREs employing maximum likelihood estimation (MLE) yields a time-efficient increase of the signal-to-noise ratio (SNR) and reduced noise bias. Further acquisition parameters included FOV = 126×126×104 mm^3^, TR = 10.4 s; TE = [53.5, 66, 78.5, 91] ms, BW = 1100 Hz/pixel, coronal orientation, foot-head PE direction, 18 segments, 1.14ms echo spacing. Sixty unique directions of the diffusion-weighting gradient were acquired with *b* = 5000 s/mm^2^ in six batches of 10 scans and interleaved with seven acquisitions with *b* = 0 (TA ≈ 36 h). Emphasis was given to intermediate breaks to maintain steady tissue temperature throughout the entire scanning session.

### 2.4. Image processing

The complex-valued GRE data from each coil channel (*n* = 32) were saved individually for each orientation and the channel combination (with preservation of the phase information) was performed offline, based on singular value decomposition (SVD) and ESPIRiT (iTerative Eigenvector-based Self-consistent Parallel Imaging Reconstruction) (Bilgic et al., 2016b; Metere & Möller, 2017; Uecker et al., 2021; Uecker & Lustig, 2017). The multi-orientation phase volumes were registered to the reference employing transformations that were derived from registrations of the corresponding magnitude volumes using FSL-FLIRT (FSL 5.0.9) (Jenkinson et al., 2012) with 6-parameter rigid transformations, a normalized mutual information (NMI) cost function, and spline interpolation. Phase unwrapping was performed on the registered phase volumes acquired at TE = 29.48 ms using the Laplacian method (Schofield & Zhu, 2003), background-phase removal using V-SHARP (Variable-kernel Sophisticated Harmonic Artifact Reduction for Phase data) (Li et al., 2011; Özbay et al, 2017; Schweser et al., 2011), and field-to-source inversion, individually for all orientations, using an iterative LSQR solver in Matlab (iLSQR) (Li et al., 2011; 2015). The iLSQR algorithm as implemented in the STI Suite (Li et al, 2014) was also employed for susceptibility-tensor reconstruction and a decomposition into its eigenvalues (χ_1_, χ_2_, χ_3_) and corresponding eigenvectors. Additionally, the previously introduced mean magnetic susceptibility (MMS) and magnetic susceptibility anisotropy (MSA) were calculated as orientation-independent tensor measures (Li et al., 2017). Separately, corresponding diffusion tensor reconstructions from the DWI data as well as calculations of eigenvalues (*λ*_1_, *λ*_2_, *λ*_3_), eigenvectors, MD and fractional anisotropy (FA) were performed using the DTIFIT tool of the FMRIB Diffusion Toolbox (FDT; FSL 5.0.9).

Following registration of the GRE and DWI acquisitions to the same reference, voxel-wise CSA-ODFs were estimated separately from the orientation-dependent phase or diffusivity data using the DIPY package (Python). The DTI-based FA metric was employed for deriving a mask including only regions of sufficient tissue anisotropy (FA > 0.2). Further processing of the local phase data included outlier exclusion by thresholding and normalization to [0,1] prior to the ODF estimation (Figure 2).

**Figure 2.**
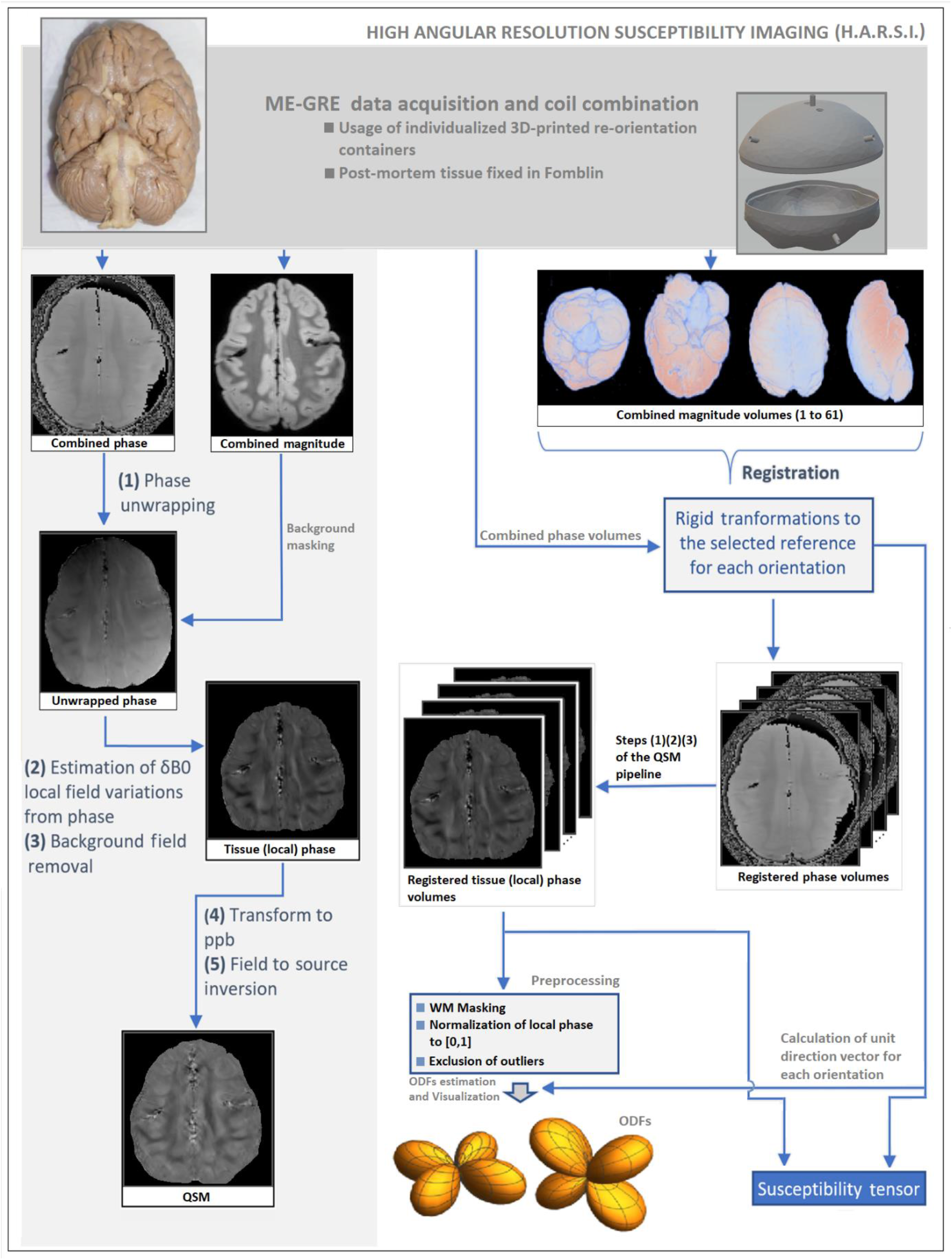
Simplified schematics of the pipelines employed for QSM as well as STI and ODF derivation from high-angular GRE phase data. The complex-valued GRE data from each of the 32 coil channels are saved individually and combined using SVD-ESPIRiT. **(Left)** QSM pipeline starting with Laplacian phase unwrapping on the combined raw phase and local *δB*_0_ estimation and removal using V-SHARP and, finally, field-to-source inversion, to obtain relative χ values employing iLSQR. **(Right)** STI and HARSI pipeline with registration of the multi-orientation phase volumes to a reference employing transformations derived from registrations of the corresponding magnitude volumes. Phase unwrapping and background-phase removal is performed individually for each orientation. iLSQR is also employed for susceptibility-tensor reconstruction. For ODF estimation, further pre-processing (masking, normalization, outlier removal) is required.

For quality assessment, the SNR was calculated for each registered GRE volume based on the magnitude images. Briefly, GM and WM masks were calculated using thresholding on the reference GRE dataset, and a 3D region of interest (ROI) was defined within ranges of ±20 voxels in anterior-posterior, ±50 voxels in inferior-superior and ±25 voxels in right-left direction around the center (Figure 3A–C). The ROI included the most significant WM areas as well as the cortical and subcortical GM. After isolating the ROI and masking, all remaining voxels were included in the analysis. Finally, an ROI of 30×30×30 voxels was identified within the artifact-free background to assess the noise, and simplified SNR estimates in GM and WM were obtained from the average signal intensities in the tissue segment divided by the standard deviation (SD) of the noise.

**Figure 3.**
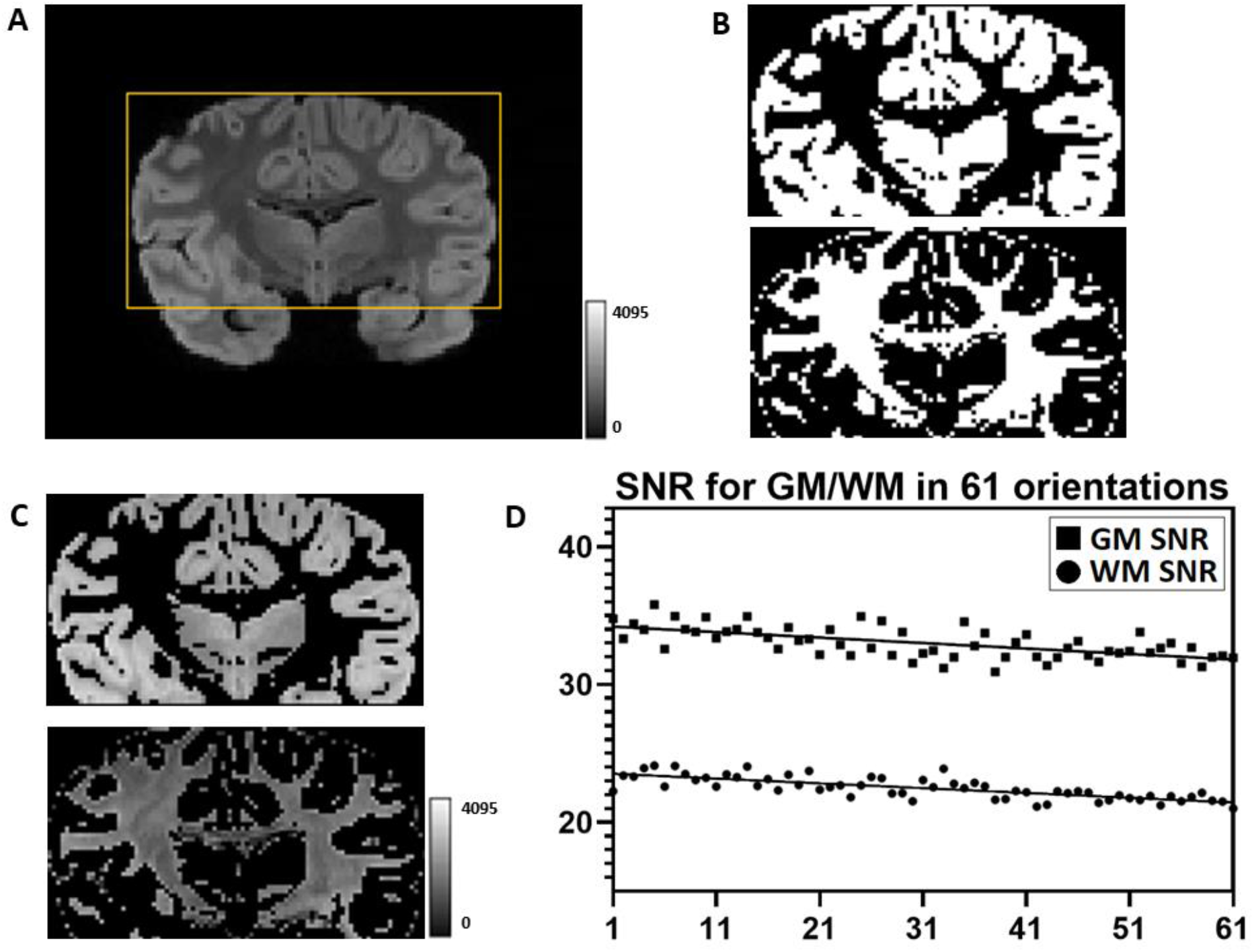
**(A)** Indication of the ROI (central coronal slice) used for the SNR calculation of the magnitude GRE images and **(B)** GM and WM masks derived by thresholding of the reference image acquired at TE=29.48 ms as well as **(C)** corresponding (magnitude) signal intensities in these masks. **(D)** The SNR metrics obtained from the 61 registered, consecutively reoriented magnitude volumes shows minor fluctuations about the reference value in both WM (circles) and GM (squares) as well as a subtle drift. Note that the orientations in (D) are ordered according to the time of the individual acquisitions. The drifts can be fitted to straight lines, which were separately calculated for the WM and GM segments.

## 3. Results

### 3.1. Data quality

The image quality of the ME-GRE acquisitions at different orientations is demonstrated in Figure 4A. Visual assessment of the local (tissue) phase of the reference volume for identifying artifacts indicated the presence of multiple small air bubbles within the cavity of the left lateral ventricle as well as single bubbles in two other regions (Figure 4B). This verified that the bubble-removal procedure was successful in most parts of the specimen and that remaining artifacts did not degrade the data quality in regions selected for the further analysis. Due to the use of the close-fitting anatomically shaped container, FSL-FLIRT achieved consistent registrations to the reference of both the orientation-dependent GRE data as well as the DWI data. Maximum deviations in the SNR of the orientation-dependent GRE acquisitions were within ±8.4% compared to the reference result indicating consistent quality throughout the experiment (mean SNR±SD: 33.0±1.1, range: 30.9–35.8 in GM and mean SNR±SD: 22.5±0.8, range: 20.9–23.9 in WM). Figure 3D further indicates an approximately linear decay of the SNR in both segments as a function of time. This decay during more than 12h scan time is of similar magnitude as the scan-to-scan SNR fluctuations and probably related to subtle drifts of the main field and shim currents. As such drift effects are corrected by the background-phase removal of the QSM pipeline, they do not lead to a relevant degradation of the phase data. Visual inspection of the DWI data (Figure 4C) indicates that an excellent quality is achieved with the segmented ME sequence implementation and the Connectom gradients, despite the substantially reduced diffusivity in fixed tissue at room temperature.

**Figure 4.**
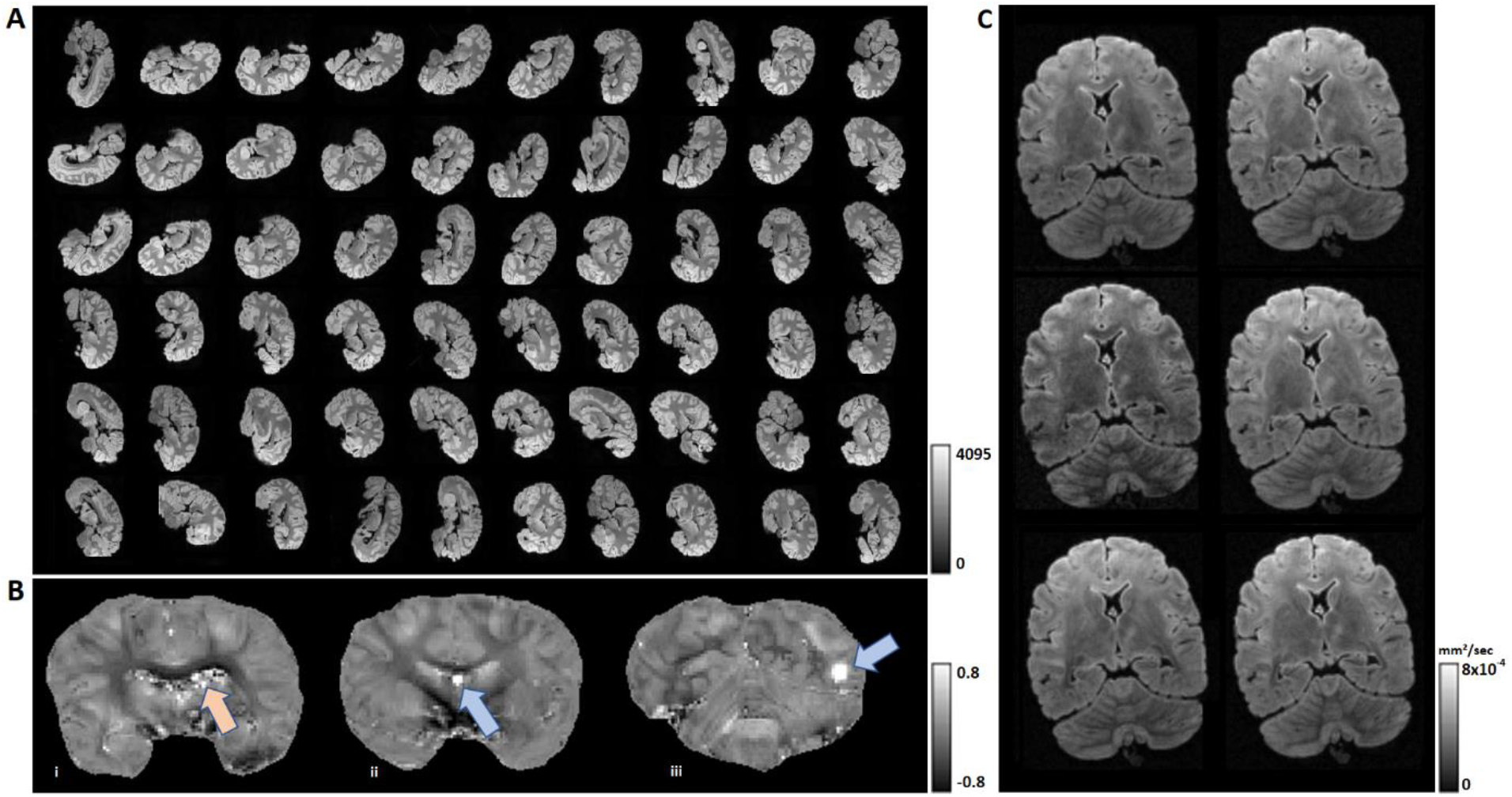
**(A)** Demonstration of the magnitude image quality obtained at TE=29.48 ms in 61 consecutive GRE acquisitions with reorientation of the sample (center slice of each individual dataset). **(B)** Local phase images at the reference orientation indicating artifacts related to remaining small air bubbles within the cavity of the left lateral ventricle (left; orange arrow) as well as at two other positions (middle and right; blue arrows). **(C)** Examples of images obtained at different directions of the diffusion-weighting gradient for demonstration of the data quality of the DWI experiment.

### 3.2. Tensor-based analyses

The high-angular GRE phase dataset provided unprecedented susceptibility tensor quality (Figure 5A). Similarly, an excellent diffusion-tensor quality was obtained with DWI, which was of equivalent spatial and angular resolution as the GRE acquisitions (Figure 5B). Consistently, the eigen-analysis of the diffusion data yielded a high quality of the MD and FA. The corresponding susceptibility metrics, MMS and MSA, yielded robust differentiation between WM and GM. The MMS exhibited local differences also within WM, indicating a particular sensitivity to the underlying microstructure of the voxels. The MSA results suggested a high sensitivity to microstructural changes but also to spurious noise from residual artifacts. Due to this slightly enhanced noise sensitivity of MSA, the DTI-derived FA map was selected as a more robust indicator of anisotropy.

**Figure 5.**
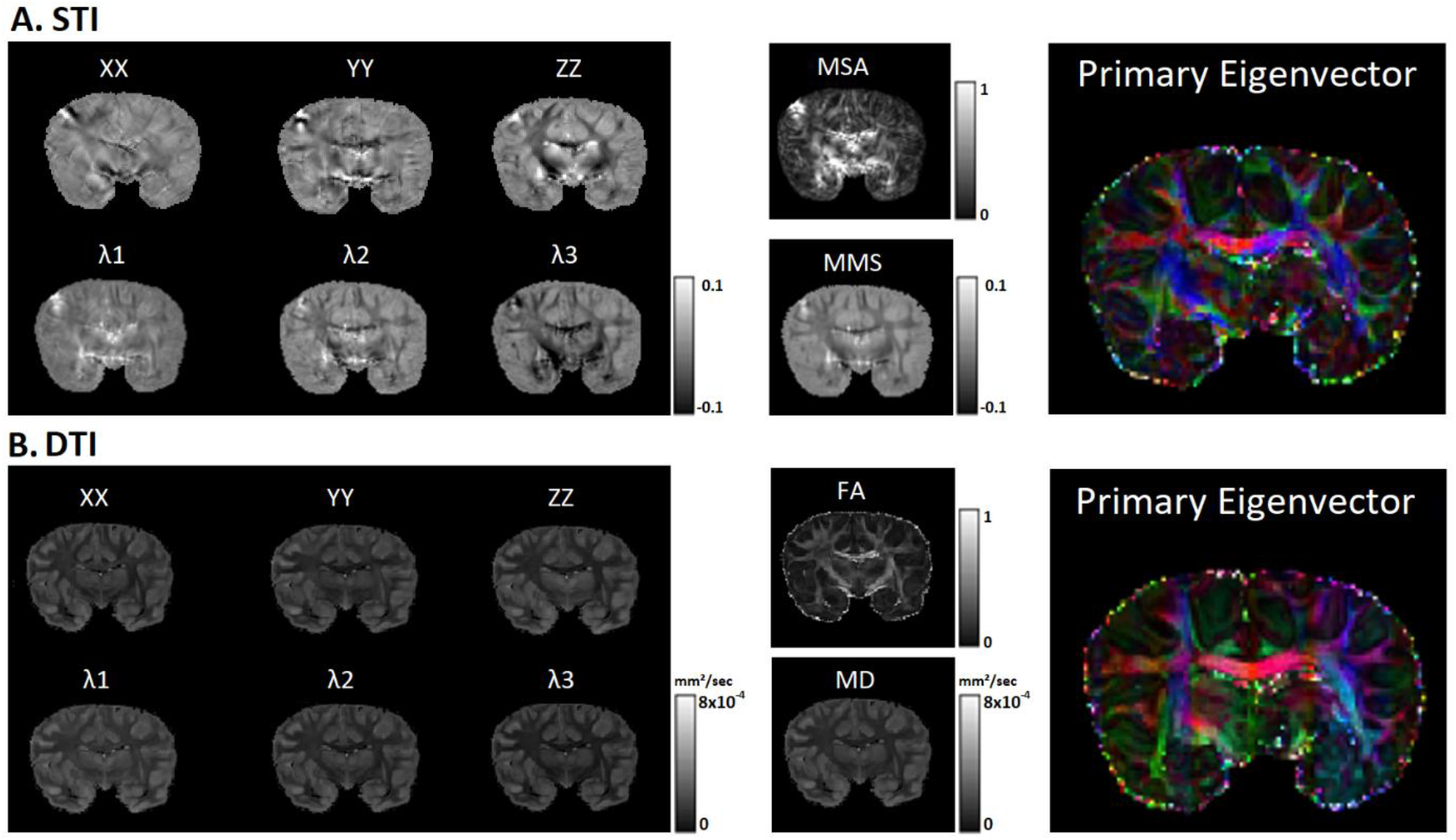
**(A)** Diagonal susceptibility-tensor components obtained with STI (left column, top row) and corresponding eigenvalues (left column, bottom row) as well as MMA and MSS (middle column) and color-coded susceptibility primary eigenvector weighted by the DTI-based FA (right column). **(B)** Diagonal diffusion-tensor components obtained with DTI (left column, top row) and corresponding eigenvalues (left column, bottom row) as well as FA and MD (middle column) and color-coded diffusivity primary eigenvector weighted by the DTI-based FA (right column). Note resemblances but also differences between the directions of the primary eigenvectors obtained with STI and DTI.

The similarity of features extracted within WM with STI and DTI as indicated by a comparison of the color-coded primary eigenvectors in Figure 5 (weighted by the FA for better visualization) is consistent with previous results employing a more restricted variation of orientations (Li et al., 2017). Note that the limitations inherent to mono-tensorial approaches allow for the identification of only a single maximum per voxel, corresponding to an average main fiber direction. Given this limitation, a qualitative resemblance of the STI and DTI results is obvious. Absolute similarity may not be expected due to the multi-step post-processing pipeline required for STI as well as contributions to susceptibility from other sources (e.g., iron) besides myelinated axons and differential sensitivities to fiber crossing between STI and DTI.

### 3.3. Impact from anisotropy in QSM

The ‘scalar’ susceptibility reference derived from STI (i.e., the χ_33_ tensor component) and from QSM obtained under the assumption of a scalar isotropic susceptibility for 10 exemplary orientations (of the total of 61 acquisitions) are presented in Figure 6A. Visual inspection yields obvious differences between acquisitions obtained with different orientation of the specimen in the magnetic field. As expected, these differences are pronounced in WM regions, where the local anisotropy is high. The absolute difference between the reference χ_33_ and the estimated χ obtained with acquisitions at different specimen orientation was investigated quantitatively in three selected WM ROIs (Figure 6B), yielding a fluctuation between 0.01 and 0.045 ppm (mean: 0.022 ppm, median: 0.021 ppm). Among these regions, those with higher structural anisotropy yielded larger differences upon reorienting the specimen. The corpus callosum was characterized by the largest range of variations, followed a similar albeit reduced effect obtained in other selected WM regions. The structural similarity index measure (SSIM) between χ_33_ and QSM at different orientations within the entire volume (Figure 6C) fluctuated between 78 and 83% (mean: 81%, median: 81%). Additional voxel-wise mapping of the variance of differences between the STI-based χ_33_ reference and QSM estimations at different orientations also indicated increased variance in WM areas associated with higher anisotropy (Figure 6D). Such variance variations were observed even within bundles. These resemble previously reported comparisons of QSM and STI-based reference results underlining the need for considering orientation dependence in WM χ estimations targeted at high accuracy.

**Figure 6.**
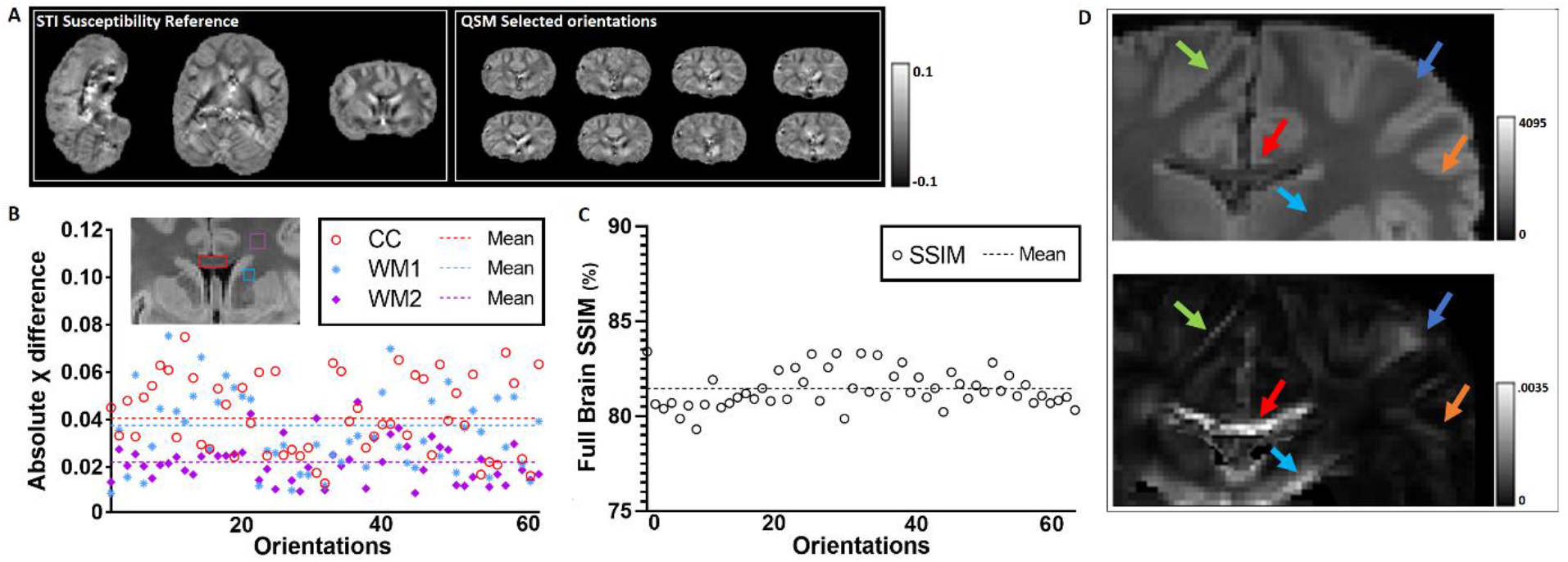
QSM analyses performed under the assumption of a scalar susceptibility. **(A)** STI-based results taking the χ_33_ tensor component (Langkammer et al., 2018) as a reference susceptibility **(left)** and comparison with standard QSM results of eight (out of 61) exemplarily selected measurements with reorientation of the specimen in the magnetic field **(right). (B)** Absolute difference between the reference and QSM results obtained without consideration of orientation dependence in three selected WM ROIs. Larger differences are obtained in areas of higher anisotropy (e.g., corpus callosum). **(C)** SSIM between the reference and QSM results without consideration of orientation dependence varies between 78 and 83%. **(D)** Voxel-wise variance map **(bottom)** of 61 QSM results obtained with different orientations. Similar to findings with DWI, increased variance is evident in areas of high structural anisotropy. The top image shows a magnitude image at the same slice position.

### 3.4. ODF estimations

Characteristic examples of HARDI- and HARSI-derived ODFs with 4^th^-order spherical harmonics are shown in Figure 7. Visual inspection indicates that diffusion-based ODFs point mostly towards one main fiber orientation while the phase-based ODFs show sensitivity to the secondary orientations, with the results indicating clearly separated lobes of similar size, in at least 2 directions.

**Figure 7.**
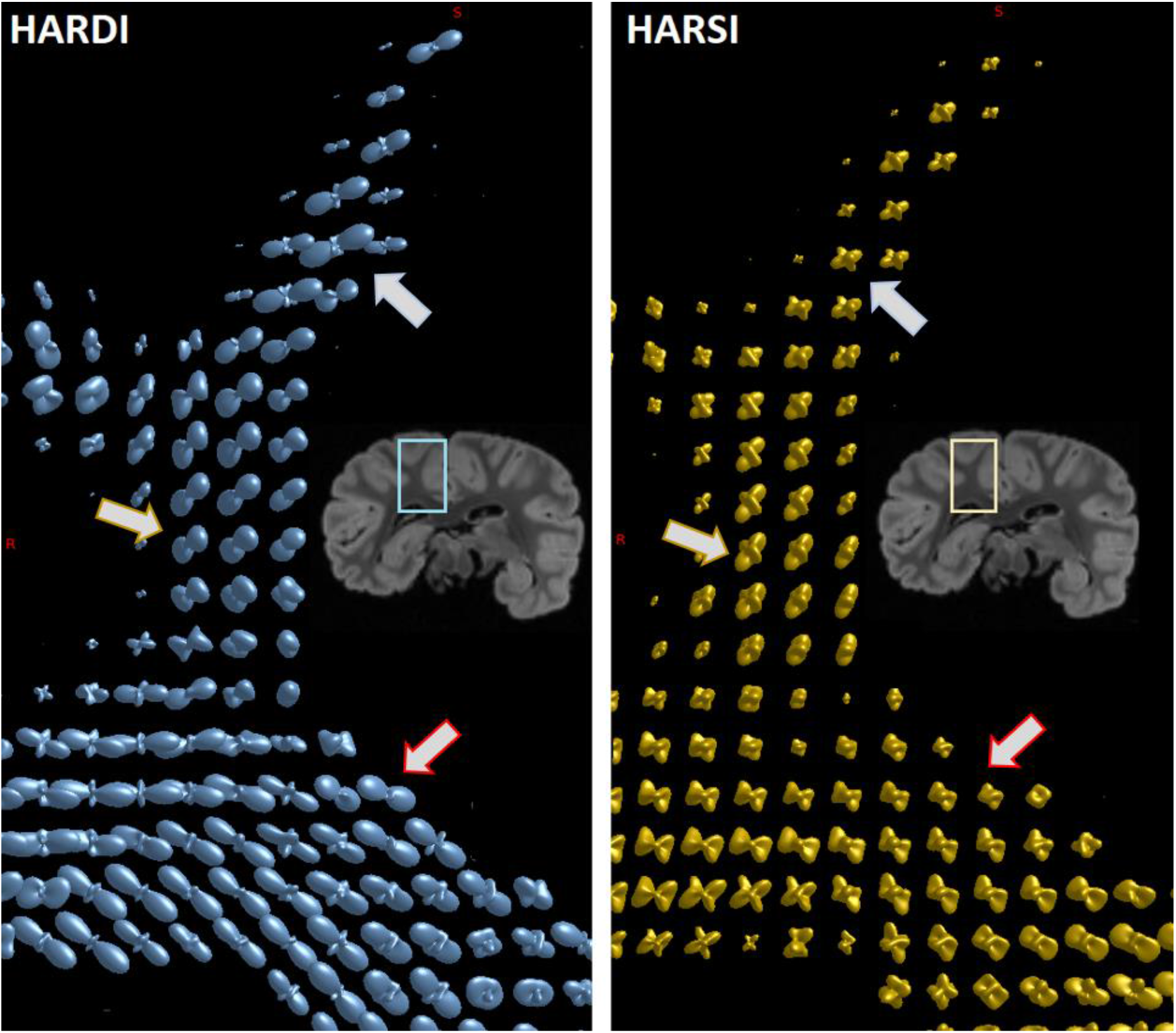
Comparison of **(A)** HARDI and **(B)** HARSI-derived ODFs in a preselected ROI with WM and surrounding GM as indicated in the insert. Color-coding of directions is not shown to better emphasize on the directionality of the estimated spherical harmonics (4^th^ order). Diffusion-based ODFs point mostly towards one main orientation, while the phase-based ODFs indicate a higher sensitivity to secondary orientations (see arrows pointing to a characteristic example).

Figure 8A, demonstrates the effect from residual air bubbles in the left lateral ventricle leading to characteristic artifacts affecting the phase-based ODFs in the surrounding region. In comparison, diffusion-based ODFs appear to be more immune to such perturbations. Further examples of ODFs obtained in artifact-free regions with diffusion and phase-based acquisitions are presented in Figure 8B–D. These results indicate an efficiency in depicting characteristics of the underlying fiber formations for both methods as well as distinct differences in the local shapes of the spherical harmonics reflecting the distributions. Further evident is some additional noise in HARSI-derived ODFs, as expected due to the multistep processing pipeline required for susceptibility data including registration of the acquisitions at different orientation.

**Figure 8.**
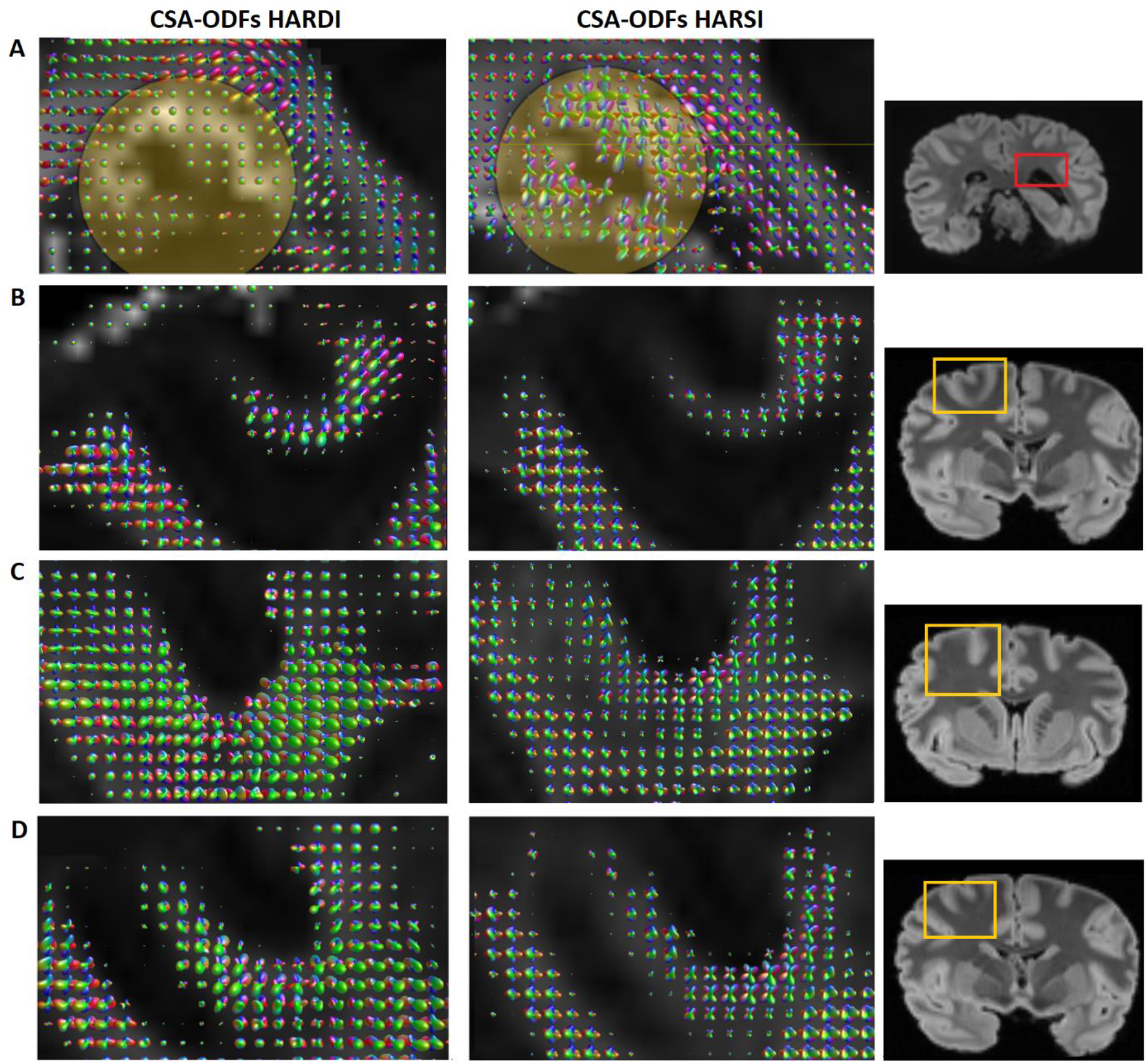
CSA-ODFs (4^th^-order spherical harmonics; FA mask as background) in selected ROIs (indicated in the right column) obtained with HARSI (left column) and HARDI (central column). **(A)** Remaining air bubbles in the left ventricle produce local field distortions leading to characteristic perturbations of the phase-based ODFs in the surrounding region (circled area), whereas the diffusion-based ODFs are relatively immune to this artifact. **(B–D)** Resemblance between the methods as well as characteristic differences in the sensitivity to second-order fibers in regions that are free from such artifacts.

## 4. Discussion

The current work goes beyond earlier investigations of orientation dependence of magnetic susceptibility in the brain in several ways: *(i)* While previous experiments largely focused on fixed rodent brain, the chimpanzee brain is morphologically much closer to the human brain with similar cellular composition and relative volume of WM (Herculano-Houzel, 2012). However, the smaller size (approx. 400 g compared to a typical human brain of 1.5 kg) allows for unconstrained reorientation in a standard head coil and more efficient sampling supporting high angular and spatial resolution acquisitions within an acceptable time. *(ii)* With 61 orientations, the angular resolution matches current DWI protocols that support extraction of information on fiber orientation distributions to identify fiber crossings or other complex patterns in WM. This permits to go beyond previous work restricted to STI. *(iii)* Simultaneously acquired DWI data of the same angular and spatial resolution support direct voxel-level comparisons of phase and diffusion-based results.

Visual comparison of the ME acquisitions at different orientations to the reference scan indicate consistent quality, which is further supported by the comparison of the achieved image SNR. Apart from local artifacts due to a few remaining air bubbles, there were no extended susceptibility artifacts, while a high quality of the registration permitted voxel-wise analysis.

The HARSI-derived ODFs indicated sensitivity to complex geometries associated with intersecting axonal fiber bundles—similar to HARDI-derived results obtained at the same spatial and angular resolution. Interestingly, a closer qualitative voxel-wise comparison of estimated fiber densities suggests—in several instances—that HARSI-based results indicate separated lobes of similar size in two main directions, whereas HARDI-based ODFs appear to point more towards a main direction, with smaller sized lobes existing towards other directions. It is well documented that the water diffusion coefficient (at ambient temperature) is reduced to 30–50% of the *in-vivo* value (at body temperature) after *in-situ* perfusion fixation (Sun et al., 2003), with further alterations occurring during the *post-mortem* interval (i.e., the interval between death and fixation) (D’Arceuil & de Crespigny, 2007; Miller et al., 2011) and during fixation (Georgi et al., 2019; Yong-Hing et al., 2005). Diffusion anisotropy reductions in WM have also been observed (D’Arceuil & de Crespigny, 2007; Miller et al., 2011), which may be related to increased membrane permeability (Shepherd et al., 2009). Aldehyde fixation finally induces moderate (*T*_1_) or strong (*T*_2_) shortening of tissue water relaxation times (Pfefferbaum et al., 2004), which is, however, mitigated by soaking the specimen in PBS (Shepherd et al., 2009) as performed in the current study. Taken together, this leads to unfavorable conditions for high-resolution DWI as correspondingly stronger *b*-values are required for resolving multiple fiber populations, without excessive prolongation of TE. For *in-vivo* MRI of human brain, *b*-values around 3,000 s/mm^2^ were recommended for an improved identification of peaks in the ODF (Jones et al, 2013). Considering the reduced diffusivity under conditions of the current work, the *b*-value of 5,000 s/mm^2^ obtained at TE = 53.5 ms with the Connectom gradient system should suffice to resolve bundles intersecting at angles ≥45° but may fall short to robustly discriminate them in the range of 30–45° (Descoteaux et al., 2009). Consequently, the HARDI-based ODFs will be increasingly dominated by contributions from one major bundle for intersections at angles <45°.

For both HARDI- and HARSI-based ODF estimations, there is no immediate information about the relative position of different fiber populations within a given voxel, which limits the differentiation of ‘crossing’ and ‘kissing’ fiber configurations (Jones et al., 2013). Minimizing the number of voxels containing multiple bundles by increasing the spatial resolution is, therefore, of ongoing interest. Diffusion-based tractography in fixed mouse brain at (isotropic) 43 µm has been achieved in previous work employing a dedicated small-bore scanner (Calabrese et al., 2015). Scanning of hominid whole-brain specimens requires large magnet bores and is technically more challenging, but also particularly interesting due to their more complex WM architecture. Resolutions around 500 µm seem to be the current limit with available hardware in such experiments (Eichner et al., 2020; Fan et al., 2022). In comparison, QSM acquisitions are less demanding on the hardware, and the signal loss at TEs in the order of 25 ms at 3 T or 12 ms at 7 T that are required for sufficient phase evolution is smaller than that inherent to DWI. Consequently, human whole-brain (magnitude) GRE datasets at 100– 200 µm are available as digital resources (Alkemade et al., 2022; Ding et al., 2016; Edlow et al., 2019). Although the precise registration of multi-orientation phase data remains challenging, the HARSI approach introduced here as a proof of concept holds great potential to go beyond current (spatial) resolution limits of diffusion-based investigations of connectivity patterns. Currently existing histological methodologies to obtain 3D information on the fiber architecture require complicated preparations (Morawski et al., 2018). These are important for a validation of MRI-derived results but currently limited to rather small brain sections.

A fundamental difference between the HARSI approach in comparison to HARDI techniques lies in the biophysical underpinning of the contrast mechanism, which is unrelated to water diffusion. In WM, myelin is a major barrier to water diffusion and contributes to diffusion anisotropy, however, diffusion anisotropy is also observed when no myelin is present (Beaulieu, 2002; Sen & Basser, 2005). Myelin also provides a primary contribution to susceptibility, however, significant susceptibility anisotropy was not observed before myelination sets in (Argyridis et al., 2014). Moreover, the signal phase is also strongly affected by the presence of paramagnetic compounds, in particular, iron stores in oligodendrocytes and astrocytes (Möller et al., 2019). Glial cells are known to cluster in short rows parallel to the axons they support (Baumann & Pham-Dinh, 2001; Suzuki & Raisman, 1992), which has recently allowed to obtain information on WM fiber architecture from Nissl stainings of *post-mortem* histological slices (Schurr & Mezer, 2021). In summary, both contributions to susceptibility in WM are roughly characterized by cylindrical geometries (hollow cylinders and rows) and should, hence, report on the particular arrangement of fibers within a voxel. A fundamental difference, however, is that the diamagnetic contribution to (anisotropic) susceptibility from myelin results from components of anisotropic molecular structures in the lipid bilayers forming myelin (Wharton & Bowtell, 2012) whereas the paramagnetic contribution may be better described as a microscopic (i.e., cellular) compartmentalization of susceptibility sources (Chu et al., 1990; He & Yablonskiy, 2009; Lee et al., 2010).

We note that the 12 echoes acquired for each orientation provide a potential for 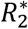 reconstruction and further analyses related to tissue structure and composition, which is beyond the scope of this study. Recently, progress has been made in separating contributions to (isotropic) QSM from opposite susceptibility sources by modeling the 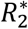 decay with multiple complex exponentials (Chen et al., 2021) or by considering the combined effects on the frequency shift and transverse relaxation rate (Shin et al., 2021). These techniques are, in general, compatible with our HARSI approach, and a corresponding combination might expand the information about microstructural tissue characteristics as well as molecular and microscopic sources of orientation dependence that are complementary to the information accessible through DWI.

## Abbreviations

3D: three-dimensional
CAD: computer aided design
CITES: Convention on International Trade in Endangered Species of Wild Fauna and Flora
CSA: constant solid angle
DTI: diffusion tensor imaging
DWI: diffusion-weighted imaging
EPI: echo planar imaging
ESPIRiT: iTerative Eigenvector-based Self-consistent Parallel Imaging Reconstruction
FDT: FMRIB Diffusion Toolbox
FLASH: Fast Low-Angle SHot
FLIRT: FMRIB’s Linear Image Registration Tool
FSL: FMRIB Software Library
GM: gray matter
GRAPPA: GeneRalized Autocalibrating Partially Parallel Acquisitions
GRE: gradient-recalled echo
HARDI: high angular resolution diffusion imaging
HARSI: high angular resolution susceptibility imaging
iLSQR: iterative LSQR
ME: multi echo
MLE: maximum likelihood estimation
MRI: magnetic resonance imaging
MP2RAGE: Magnetization-Prepared 2 RApid Gradient Echoes
NMI: normalized mutual information
ODF: orientation distribution function
PBS: phosphate-buffered saline
PE: phase-encoding
PFA: paraformaldehyde
QSM: quantitative susceptibility mapping
RF: radiofrequency
ROI: region of interest
SSIM: structural similarity index measure
STI: susceptibility tensor imaging
STL: stereolithography
SVD: singular value decomposition
V-SHARP: Variable-kernel Sophisticated Harmonic Artifact Reduction for Phase data
WM: white matter.

## Mathematical Symbols

*δB*_0_: local offset of the amplitude of the magnetic flux density;

BW: bandwidth;

*b*: *b*-value;

FA: fractional anisotropy;

FOV: field of view;

*f*_*p*_: partial-Fourier factor;

*H*_0_: magnitude of the applied magnetic field;

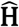: unit vector along the direction of the applied magnetic field;

**k**: spatial frequency vector;

MD: mean diffusivity;

MMS: mean magnetic susceptibility;

MSA: magnetic susceptibility anisotropy;

*n*: number (integer);

*R*: GRAPPA acceleration factor;

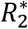: effective transverse relaxation rate;

**r**: position vector;

SD: standard deviation;

SNR: signal-to-noise ratio;

*T*_1_: longitudinal relaxation time;

TA: acquisition time;

TE: echo time;

ΔTE: echo spacing;

TR: repetition time;

*x, y, z*: cartesian coordinates;

*α*: RF pulse flip angle,

*γ*: gyromagnetic ratio;

*φ*: signal phase;

*λ*_1_, *λ*_2_, *λ*_3_: eigenvalues of the diffusion tensor;

µ_0_: vacuum permeability;

χ: bulk volume magnetic susceptibility;

χ magnetic susceptibility tensor;

χ_1_, χ_2_, χ_3_: eigenvalues of the magnetic susceptibility tensor;

χ_*ij*_: magnetic susceptibility tensor element;

ℱ, ℱ^−1^: Fourier transform and inverse Fourier transform;

^*T*^: transpose of a matrix.

## CRediT Statement

**Dimitrios G. Gkotsoulias:** Conceptualization, Methodology, Software, Validation, Formal analysis, Investigation, Writing – Original Draft, Writing - Review Editing, Visualization

**Roland Müller:** Methodology, Resources

**Carsten Jäger:** Methodology, Resources

**Torsten Schlumm:** Software, Data Curation

**Toralf Mildner:** Methodology, Resources, Writing - Review Editing

**Cornelius Eichner:** Conceptualization, Writing - Review Editing

**André Pampel:** Conceptualization, Writing - Review Editing

**Jennifer Jaffe:** Resources

**Tobias Gräßle:** Resources

**Niklas Alsleben:** Software

**Jingjia Chen:** Validation

**Catherine Crockford:** Resources, Writing - Review Editing

**Roman Wittig:** Resources, Writing - Review Editing

**Chunlei Liu:** Conceptualization, Writing - Review Editing, Funding acquisition

**Harald E. Möller:** Conceptualization, Methodology, Resources, Writing – Original Draft, Writing - Review Editing, Supervision, Project administration, Funding acquisition, Supervision

## Acknowledgements

This work was funded by the EU through the ITN “INSPiRE-MED” (H2020-MSCA-ITN-2018, #813120). Chunlei Liu and Jingjia Chen were supported in part by the National Institute of Aging of the National Institutes of Health (Award No. R01AG070826). We particularly thank Angela D. Friederici and Nikolaus Weiskopf and further Evolution of Brain Connectivity (EBC) project organizers, the Ministère de l’Enseignement Supérieur et de la Recherche Scientifique and the Ministère des Eaux et Fôrets, Côte d’Ivoire, the Office Ivoirien des Parcs et Réserves, and the staff of the Taï Chimpanzee Project for permitting and supporting this research. Appreciation is extended to Michael Paquette, Riccardo Metere and Alfred Anwander for helpful methodological discussions and sharing of related processing packages.

